# HIV and FIV glycoproteins increase cellular tau pathology via cGMP-dependent kinase II activation

**DOI:** 10.1101/2022.01.11.475910

**Authors:** Matheus F. Sathler, Michael J. Doolittle, James A. Cockrell, India R. Nadalin, Franz Hofmann, Sue VandeWoude, Seonil Kim

## Abstract

As the development of combination antiretroviral therapy (cART) against human immunodeficiency virus (HIV) drastically improves the lifespan of individuals with HIV, many are now entering the prime age when Alzheimer’s disease (AD)-like symptoms begin to manifest. Hyperphosphorylated tau, a known AD pathological characteristic, has been prematurely increased in the brains of HIV-infected patients as early as in their 30s and is increased with age. This thus suggests that HIV infection may lead to accelerated AD phenotypes. However, whether HIV infection causes AD to develop more quickly in the brain is not yet fully determined. Interestingly, we have previously revealed that viral glycoproteins, HIV gp120 and feline immunodeficiency virus (FIV) gp95, induce neuronal hyperexcitation via cGMP-dependent kinase II (cGKII) activation in cultured hippocampal neurons. Here, we use cultured mouse cortical neurons to demonstrate that HIV gp120 and FIV gp95 are sufficient to increase cellular tau pathology, including intracellular tau hyperphosphorylation and tau release to the extracellular space. We further reveal that viral glycoprotein-induced cellular tau pathology requires cGKII activation. Together, HIV infection likely accelerates AD-related tau pathology via cGKII activation.

## Introduction

Human immunodeficiency virus (HIV) continues to be a major public health issue because HIV infection has become a chronic disease with the advent of combination antiretroviral therapy (cART) that has allowed individuals infected at a younger age to survive into older age (Farhadian et al., 2017). Importantly, a large proportion of HIV patient population faces aging-associated brain disorders, including Alzheimer’s disease (AD) (Miller et al., 2011). In fact, prevalence of AD is elevated among patients with HIV and even higher among patients who are not being treated with cART (Alisky, 2007; Brew et al., 2005; Nebuloni et al., 2001). Significantly, hyperphosphorylated tau, a known AD pathological characteristic, has been prematurely increased in the brains of HIV-infected patients as early as in their 30s and is increased with age (Anthony et al., 2006). This thus suggests that HIV may lead to accelerated AD-associated tau phenotypes. However, whether HIV infection can lead to accelerated AD development in the brain remains unclear (Jha et al., 2020).

One of the major limitations in searching for mechanisms and treatments for AD in HIV infection is the lack of reliable animal models for investigating the pathophysiology (Chambers et al., 2015). Rodent models are heavily used for the study of HIV-associated neuropathology (Jaeger and Nath, 2012). However, results obtained in these models are often not easily translated to human pathology, given that rodents are not naturally susceptible to HIV infection and thus do not reflect chronic *in vivo* nature of infection (Fox and Gendelman, 2012). Although non-human primates infected with simian immunodeficiency virus (SIV) or genetic chimeras of SIV and HIV have a number of important advantages over small-animal models, they have obvious disadvantages in terms of high maintenance costs and considerable genetic variation that greatly complicate studies, especially when the number of animals used is low (Hatziioannou and Evans, 2012). Moreover, SIV is unable to cause acquired immune deficiency syndrome (AIDS) in its natural hosts (sooty mangabeys and chimpanzees), whereas SIV shows a relatively high pathogenic potential in rhesus macaques and humans following cross-species transmission events (Apetrei et al., 2005; Gardner, 1996; Sharp et al., 2005), which have extremely unbalanced epidemiologic consequences (Apetrei et al., 2005; Gardner, 1996). Most importantly, these animal models are unable to naturally develop neurofibrillary tangles (NTFs), a known tau pathology in AD (Chambers et al., 2015). Thus, new animal models to examine HIV-associated tau pathology are an important current and future need.

Feline immunodeficiency virus (FIV) infection in domestic cats represents an animal model of immunodeficiency that shares similarities in pathogenesis of HIV in humans (Meeker and Hudson, 2017). In addition, certain strains of FIV can enter the central nervous system (CNS), underlying neurological symptoms similar to those observed in some individuals infected with HIV (Elder et al., 2010). Several neurological deficits have been also found as early as 12-months post-infection in studies of experimentally infected specific pathogen free (SPF) cats (younger than 6 months) designed to mimic HIV infection in humans (Meeker and Hudson, 2017). Beta-amyloid-associated senile plaques are a major pathological hallmark in AD and are found in many animal species including chimpanzees and dogs, however, many of these animals are unable to exhibit NFTs and subsequent neurodegeneration (Chambers et al., 2015). Notably, cats express multiple tau isoforms like humans, and aged cats older than 14 years are a unique animal species that naturally replicates NFTs like humans (Chambers et al., 2015; Head et al., 2005; Janke et al., 1999). In addition, the combination of HIV antiretroviral drugs on cats naturally infected with FIV in the late phase of the asymptomatic state of the disease significantly reduces viral loads (Gomez et al., 2012). Therefore, FIV infection in domestic cats can be an important model to substantially improve our understanding the HIV-induced tau pathophysiology relevant to an older population of HIV-infected patients with antiretroviral therapy.

As the development of cART against HIV drastically reduces the number of deaths from HIV-related causes, more of individuals with HIV are now reaching older ages when AD-like symptoms are more prevalent (Chakradhar, 2018). Importantly, HIV is believed to prematurely age the brains of those with the disease and lead to brain dysfunction, as can AD (Chakradhar, 2018). Given these commonalities, it is possible that HIV could create conditions ripe for the development of AD (Chakradhar, 2018). Although the molecular and cellular mechanisms underlying this connection have not been fully investigated (Chakradhar, 2018). One potential mechanism of a HIV–AD connection is excitotoxicity, the pathological process by which neurons are damaged by neuronal hyperexcitation (Hinkin et al., 1995; Rottenberg et al., 1996; von Giesen et al., 2000). However, HIV does not directly infect neurons but instead use a noninfectious interaction between the viral envelope and the neuronal surface (Bragg et al., 1999; Brenneman et al., 1988). HIV gp120 indirectly and/or directly interacts with neurons, which enhances excitatory synaptic receptor activity, resulting in synaptic damages, while the mechanisms are not currently understood (Kettenmann et al., 2013; Kim et al., 2011; Xu et al., 2011). We have previously demonstrated that both HIV gp120 and FIV gp95 interact with the receptor for the α-chemokine stromal cell-derived factor 1, CXCR4, which enhances neuronal Ca^2+^ activity through NMDA receptors (NMDARs) and activate cGMP-dependent protein kinase II (cGKII) via nitric oxide (NO) signaling (Sztukowski et al., 2018). Notably, cGKII can phosphorylate AMPA receptor (AMPAR) subunit GluA1, which triggers their synaptic trafficking, a critical step for inducing synaptic plasticity (Kim et al., 2015a; Serulle et al., 2007). Importantly, we have found that HIV gp120 and FIV gp95 induce aberrant stimulation of cGKII through the CXCR4-NMDAR-NO pathway, resulting in AMPAR-mediated Ca^2+^ hyperactivity in cultured mouse and feline hippocampal neurons (Sztukowski et al., 2018). This suggests that viral glycoproteins induce neuronal hyperexcitation via cGKII activation. This hyperexcitation is also strongly associated with early AD pathogenesis (Bero et al., 2011; Cirrito et al., 2008; Gibbons et al., 2019; Pooler et al., 2013; Wu et al., 2016; Yamada et al., 2014). Therefore, neuronal cGKII activation induced by HIV gp120 and FIV gp95 may provide a molecular link between HIV and AD.

Here, our data using cultured mouse cortical neurons have demonstrated that HIV gp120 and FIV gp95 significantly increases cellular tau pathology, including intracellular hyperphosphorylated tau and extracellular tau release. In addition, we have found that this cellular tau pathology is dependent on cGKII activation. Together, existing data and our new findings have revealed that HIV infection accelerates AD-related neural hyperexcitation and tau pathology via cGKII activation.

## Results

### HIV gp120 or FIV gp95 treatment significantly increases neuronal Ca^2+^ activity in cultured mouse cortical neurons via cGKII activation

Neuronal Ca^2+^ is the second messenger responsible for transmitting depolarization status and synaptic activity (Gleichmann and Mattson, 2011). Ca^2+^ regulation is a vital process in neurons because of these characteristics, and abnormal Ca^2+^ activity in neurons is one of the major contributors to many neurological diseases (Gleichmann and Mattson, 2011). By measuring neuronal Ca^2+^ activity, we have previously shown that HIV gp120 or FIV gp95 treatment significantly increases neuronal activity in cultured mouse and feline hippocampal neurons (Sztukowski et al., 2018). We thus examined whether HIV gp120 or FIV gp95 treatment affected Ca^2+^ activity in cultured wild-type (WT) mouse cortical neurons infected with adeno-associated virus (AAV) expressing GCaMP7s, a genetically encoded Ca^2+^ indicator (Dana et al., 2019). First, we acutely treated day *in vitro* (DIV) 12 – 14 neurons with 1 nM HIV gp120 or 1 nM FIV gp95 and determined Ca^2+^ activity immediately after treatment as described previously (Kim et al., 2015a; Kim et al., 2015b; Kim and Ziff, 2014; Roberts et al., 2021; Sun et al., 2019; Sztukowski et al., 2018). We found active spontaneous Ca^2+^ transients in control cells (CTRL) and neurons treated with viral proteins (**Fig. 1A**). However, total Ca^2+^ activity in HIV gp120 or FIV gp95-treated cells was significantly higher than in controls (CTRL, 1 ± 0.41 ΔF/F_min_; HIV gp120, 1.70 ± 0.51 ΔF/F_min_, *p* < 0.0001, and FIV gp95, 1.55 ± 0.67 ΔF/F_min_, *p* = 0.0002), confirming that viral glycoproteins were sufficient to increase neuronal Ca^2+^ activity in cultured mouse cortical neurons (**Fig. 1A**). Importantly, Ca^2+^ flux through the NMDAR-nNOS pathway activates cGKII by the production of cGMP (Bredt, 2003). cGKII mediates phosphorylation of serine 845 of GluA1 (pGluA1), important for activity-dependent trafficking of GluA1-containing AMPARs, and increases the level of extrasynaptic receptors (Kim et al., 2015a; Serulle et al., 2007). We have also shown that HIV gp120 or FIV gp95 treatment activates cGKII, which subsequently phosphorylates AMPAR subunit GluA1, leading to the elevation of surface GluA1 expression and AMPAR-mediated synaptic activity, a cellular basis of hyperexcitation in viral glycoprotein-treated cultured hippocampal neurons (Sztukowski et al., 2018). We thus tested the possibility that cGKII was a downstream effector of viral glycoprotein-induced Ca^2+^ hyperexcitation in cultured cortical neurons. We found that HIV gp120 or FIV gp95 treatment was unable to increase Ca^2+^ activity when cGKII activity was blocked by treating neurons with 1 μM Rp8-Br-PET-cGMPS (RP), a cGKII inhibitor (HIV gp120 + RP, 1.19 ± 0.51 ΔF/F_min_, *p* = 0.0005, and FIV gp95 + RP, 1.18 ± 0.64 ΔF/F_min_, *p* = 0.0388) (**Fig. 1A**). However, RP treatment in control neurons had no effect on Ca^2+^ activity (CTRL + RP, 1.04 ± 0.53 ΔF/F_min_, *p* = 0.9992) (**Fig. 1A**). We further cultured cortical neurons from cGKII knockout (KO) mice as described previously (Kim et al., 2015a; Sztukowski et al., 2018) and confirmed that 1 nM HIV gp120 or 1 nM FIV gp95 treatment had no effect on Ca^2+^ dynamics in KO neurons (CTRL, 1 ± 0.54 ΔF/F_min_, HIV gp120, 0.85 ± 0.50 ΔF/F_min_, *p* = 0.2560, and FIV gp95, 1.00 ± 0.50 ΔF/F_min_ *p* = 0.9988) (**Fig. 1B**). This suggests that viral glycoproteins, HIV gp120 and FIV gp95, share the cGKII-mediated core cellular pathway to induce neuronal hyperexcitation in cultured mouse cortical neurons, which is consistent with the previous findings in cultured mouse and feline hippocampal neurons (Sztukowski et al., 2018).

**Figure 1.**
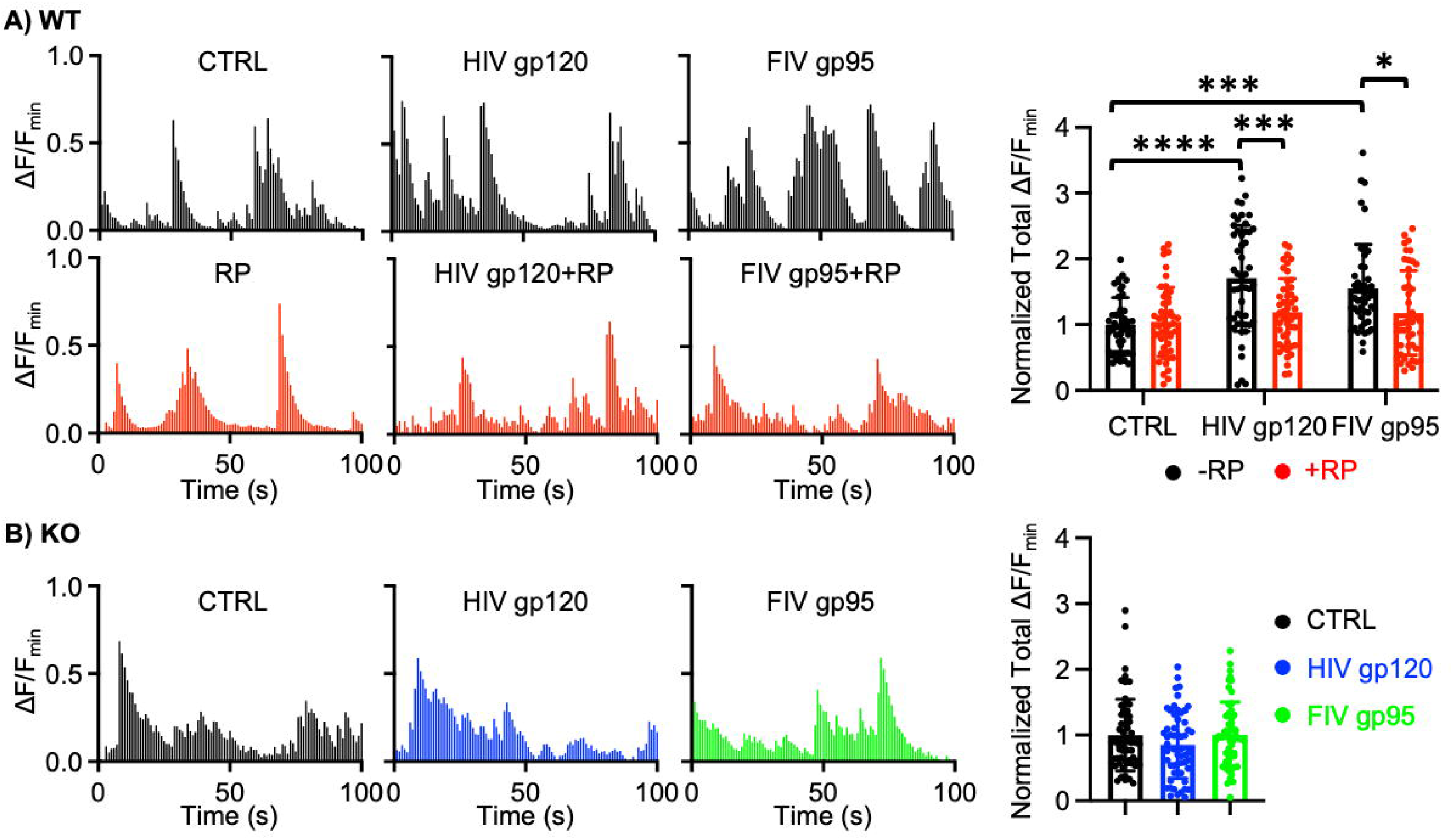
HIV gp120 or FIV gp95 treatment significantly increases neuronal Ca^2+^ activity in cultured mouse cortical neurons via cGKII activation. **A)** Representative traces of GCaMP7s fluorescence intensity and a summary graph of normalized average of total Ca^2+^ activity in cultured WT cortical neurons in each condition showing that 1 nM HIV gp120 or 1 nM FIV gp95 treatment significantly increases neuronal Ca^2+^ activity, which is reversed by inhibition of cGKII activity using 1 μM Rp8-Br-PET-cGMPS (RP) (n = number of neurons, control (CTRL) = 48, RP = 49, HIV gp120 = 50, HIV gp120 + RP = 49, FIV gp95 = 47, and FIV gp95 + RP = 46, \**p* < 0.05, \*\*\**p* < 0.001, and \*\*\*\**p* < 0.0001, Two-way ANOVA with Tukey test). **B)** Representative traces of GCaMP7s fluorescence intensity and a summary graph of normalized average of total Ca^2+^ activity in cultured cGKII KO cortical neurons in each condition showing that 1 nM HIV gp120 or 1 nM FIV gp95 treatment has no effect on neuronal Ca^2+^ activity in KO cells (n = number of neurons, control (CTRL) = 60, HIV gp120 = 56, and FIV gp95 = 56, One-way ANOVA with Tukey test).

### HIV gp120 or FIV gp95 treatment significantly increases extracellular tau levels via cGKII activation

Although tau is primarily a cytoplasmic protein that stabilizes microtubules, it can be released into the extracellular space, enter neighboring neurons, and spread tau pathology throughout the brain, which is not dependent on neuronal death but can be stimulated by increased neuronal activity as AD progresses (Braak and Braak, 1991; Chai et al., 2012; Clavaguera et al., 2009; Frost et al., 2009; Guo and Lee, 2011; Iba et al., 2013; Karch et al., 2012; Kfoury et al., 2012; Pooler et al., 2013; Sanders et al., 2014; Wu et al., 2016; Yamada et al., 2014). Several recent studies have showed that tau can be physiologically released to the extracellular fluid both *in vivo* and in cultured cells, and such release appears to be regulated by neuronal activity (Pooler et al., 2013; Wu et al., 2016; Yamada et al., 2014). We thus determined whether an increase in neuronal activity was sufficient to induce extracellular tau release in cultured mouse cortical neurons. Extracellular tau levels were measured using a total tau solid-phase sandwich enzyme-linked immunosorbent assay (ELISA) with a mouse tau antibody (Yan et al., 2016). To directly activate neurons’ activity, we treated DIV 14 neurons with 1 μM glutamate, an excitatory neurotransmitter, for 10 min and collected the conditioned culture medium to measure extracellular tau concentration. As expected, glutamate treatment significantly increased extracellular tau levels compared to control cells (CTRL, 631.45 ± 188.60 pg/ml and glutamate 1,341.52 ± 266.63 pg/ml, *p* = 0.0048) (**Fig. 2A**), confirming that elevated neuronal activity was sufficient to increase extracellular tau levels in cultured neurons. Given that HIV gp120 or FIV gp95 treatment is sufficient to increase neuronal activity in cultured mouse cortical neurons (**Fig. 1**), we next examined whether viral glycoproteins enhanced tau release to the extracellular space in cultured neurons. To test this, we treated DIV 14 neurons with 1 nM HIV gp120 or 1 nM FIV gp95 for 24 hours and measured extracellular tau levels in the conditioned culture medium by ELISA as described above. We found that both HIV gp120 and FIV gp95 significantly increased extracellular tau concentration (CTRL, 661.10 ± 169.09 pg/ml, HIV gp120, 1,506.31 ± 224.96 pg/ml, *p* = 0.0055, and FIV gp95, 987.11 ± 250.15 pg/ml, *p* = 0.0325) (**Fig. 2B**). This suggests that viral glycoprotein-induced neuronal hyperexcitation contributes to an increase in tau release to the extracellular space. Given that viral glycoprotein-induced neuronal hyperexcitation is dependent on cGKII activation (**Fig. 1**), we examined whether inhibition of cGKII activity reversed the viral glycoprotein effects on extracellular tau levels in cultured neurons. 1 μM RP was added to DIV 14 cultured neurons treated with 1 nM HIV gp120 or 1 nM FIV gp95 for 24 hours, and conditioned culture medium was collected to determine extracellular tau levels by ELISA as described above. We found that inhibition of cGKII activity significantly reversed viral glycoprotein-induced increased extracellular tau concentrations (HIV gp120 + RP, 516.17 ± 131.04 pg/ml, *p* = 0.0003, and FIV gp95 + RP, 526.82 ± 219.69 pg/ml, *p* = 0.0024) (**Fig. 2B**). However, RP treatment in the absence of viral glycoproteins had no effect on tau release (CTRL + RP, 571.35 ± 193.31 pg/ml, *p* = 0.9634) (**Fig. 2B**). To further address whether cGKII activation was crucial for an increase in extracellular tau levels by viral glycoproteins, we used cGKII KO cultured cortical neurons and treated with 1 nM HIV gp120 or 1 nM FIV gp95 for 24 hours, extracellular tau concentration was measured by ELISA as described above. We found that HIV gp120 or FIV gp95 treatment was unable to increase extracellular tau levels in cGKII KO neurons (CTRL, 588.10 ± 198.33 pg/ml, HIV gp120, 644.23 ± 272.66 pg/ml, *p* = 0.8633, and FIV gp95, 617.97 ± 313.13 pg/ml *p* = 0.9591) (**Fig. 2C**). Taken together, viral glycoproteins, HIV gp120 and FIV gp95, are sufficient to increase extracellular tau concentration via cGKII-dependent neuronal hyperexcitation in cultured cortical neurons.

**Figure 2.**
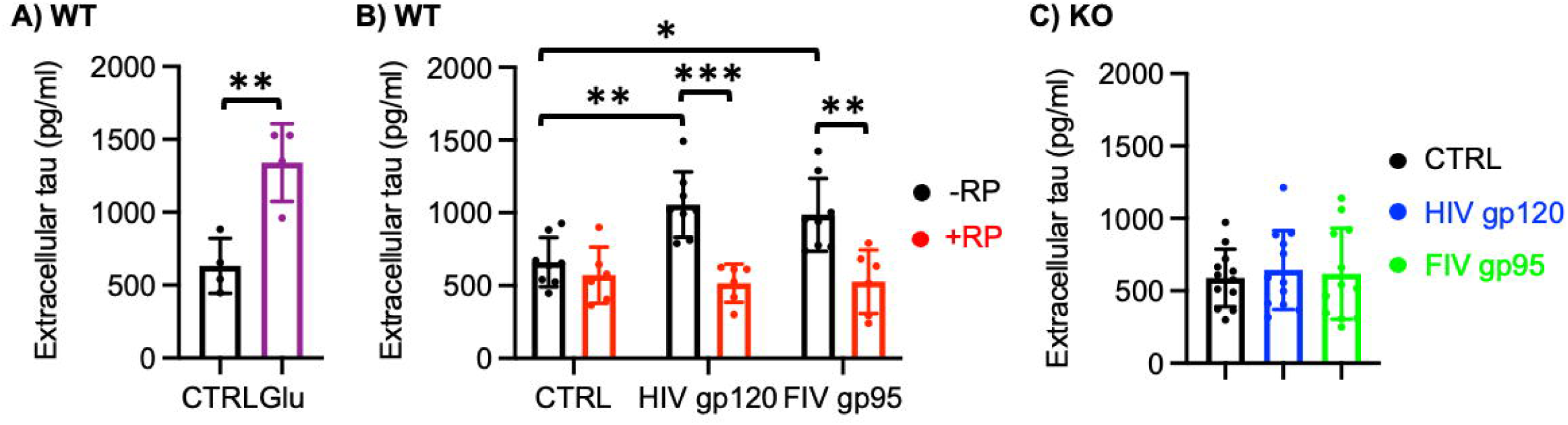
HIV gp120 or FIV gp95 treatment significantly increases extracellular tau levels via cGKII activation. **A**) A summary graph of extracellular tau levels in cultured WT cortical neurons in the absence or presence of 1 μM glutamate (Glu) showing that an increase in neuronal activity significantly increases extracellular tau concentration (n = number of ELISA measurements, control (CTRL) = 4 and Glu = 4, \**p* < 0.05, unpaired student t-test). **B**) A summary graph of extracellular tau levels in cultured WT cortical neurons in each condition demonstrating that viral glycoproteins, HIV gp120 and FIV gp95, significantly elevates tau release to the extracellular space, which is reversed by cGKII inhibition using 1 μM Rp8-Br-PET-cGMPS (RP) (n = number of ELISA measurements, control (CTRL) = 8, RP = 6, HIV gp120 = 8, HIV gp120 + RP = 6, FIV gp95 = 8, and FIV gp95 + RP = 6, \**p* < 0.05, \*\**p* < 0.01, and \*\*\**p* < 0.001, Two-way ANOVA with Tukey test). **C**) A summary graph of extracellular tau levels in cultured cGKII KO cortical neurons treated with 1 nM HIV gp120 or 1 nM FIV gp95 showing that viral glycoproteins are unable to alter extracellular tau levels in KO cells (n = number of ELISA measurements, control (CTRL) = 12, HIV gp120 = 12, and FIV gp95 = 12, One-way ANOVA with Tukey test).

### Viral glycoproteins, HIV gp120 and FIV gp95, significantly enhance tau hyperphosphorylation via cGKII activation

Hyperphosphorylated tau has been identified in the brains of HIV-positive people with brain impairment as early as their 30s, and its levels rise with age, which is known to contribute to neurodegeneration in HIV brains (Anthony et al., 2006; Patrick et al., 2011). HIV gp120 is known to induce tau hyperphosphorylation, which in turn contributes to NFT formation in rodent brains (Cho et al., 2017; Kang et al., 2010). We thus examined whether HIV gp120 or FIV gp95 treatment induced tau hyperphosphorylation in cultured mouse cortical neurons. DIV 14 cultured cortical neurons were treated with 1 nM HIV gp120 or 1 nM FIV gp95 for 24 hours, and total cell lysates were collected to determine tau hyperphosphorylation using immunoblots. We found that HIV gp120 or FIV gp95 treatment significantly increased AT8 antibody (recognizing tau phosphorylation at serine 202 and threonine 205)-positive phosphorylated paired helical filament (PHF) tau (CTRL, 1.00, HIV gp120, 1.92 ± 0.49, *p* < 0.0001, and FIV gp95, 1.47 ± 0.35, *p* = 0.0012) (**Fig. 3A**). As we have shown that cGKII activation is critical for viral glycoprotein-induced neuronal hyperexcitation (**Fig. 1**) and elevated extracellular tau levels (**Fig. 2**), we examined whether viral glycoprotein-induced cGKII stimulation was responsible for tau hyperphosphorylation. 1 μM RP was added to DIV 14 cultured neurons treated with 1 nM HIV gp120 or 1 nM FIV gp95 for 24 hours, and hyperphosphorylated tau levels were examined using immunoblots. Notably, inhibition of cGKII activation was sufficient to abolish a HIV gp120 or FIV gp95-induced increase in phosphorylated PHF tau levels (HIV gp120 + RP, 0.82 ± 0.29, *p* < 0.0001, and FIV gp95, 0.97 ± 0.28, *p* = 0.0026) (**Fig. 3A**). In contrast, RP treatment in control cells was unable to affect tau hyperphosphorylation (CTRL + RP, 1.00 ± 0.40, *p* > 0.9999) (**Fig. 3A**). To further address whether cGKII activation was important for viral glycoprotein-induced tau PHF formation, we employed cGKII KO cultured cortical neurons and treated with 1 nM HIV gp120 or 1 nM FIV gp95 for 24 hours, hyperphosphorylated tau was determined by immunoblot using the AT8 antibody as described above. We found that HIV gp120 or FIV gp95 treatment was unable to increase phosphorylated PHF tau levels in cGKII KO neurons (CTRL, 1.00, HIV gp120, 1.10 ± 0.95, *p* = 0.9841, and FIV gp95, 0.87 ± 0.46, *p* = 0.9513) (**Fig. 3B**). Taken together, viral glycoproteins, HIV gp120 and FIV gp95, are sufficient to induce tau hyperphosphorylation via cGKII activation in cultured cortical neurons.

**Figure 3.**
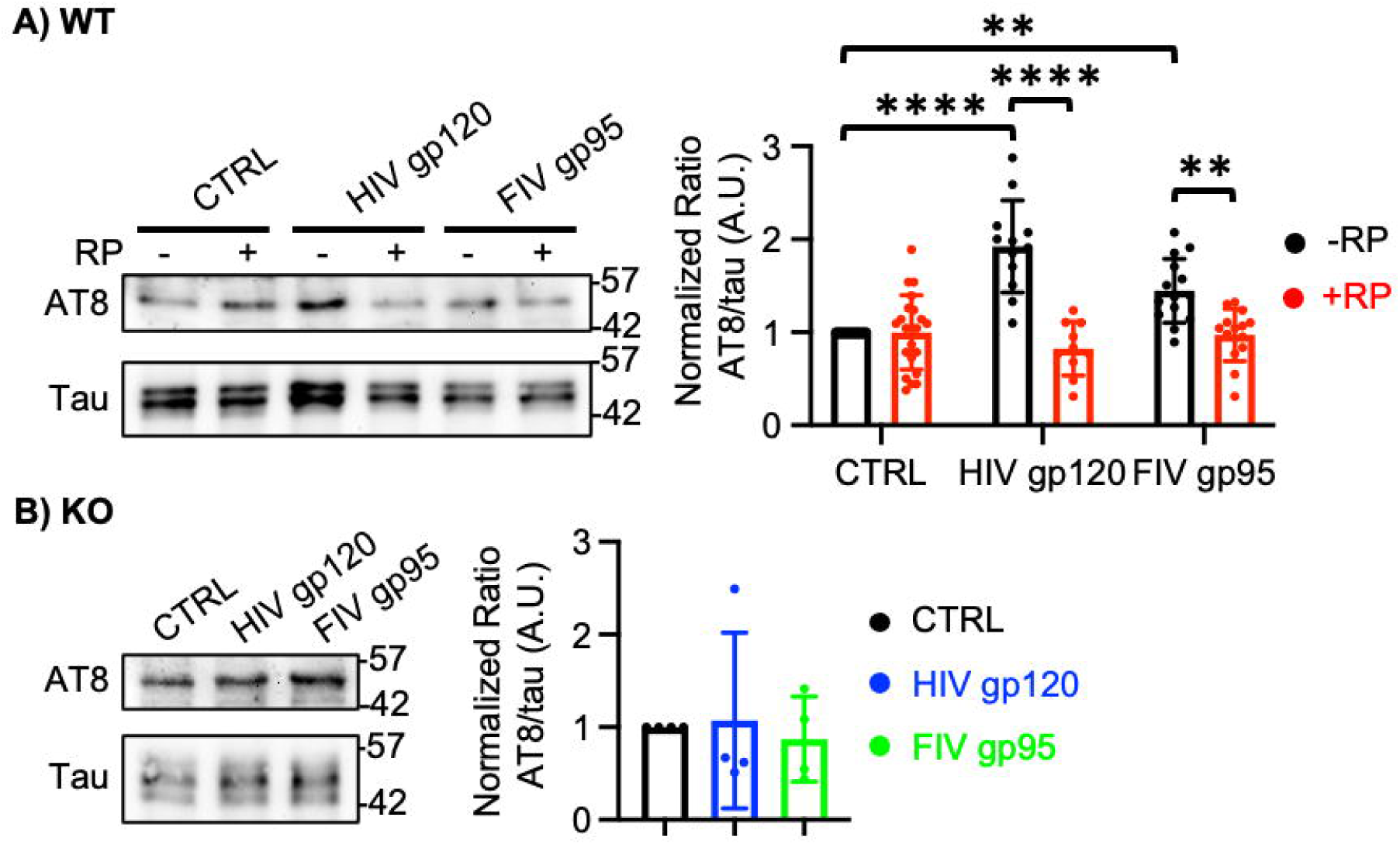
Viral glycoproteins, HIV gp120 and FIV gp95, significantly enhance tau hyperphosphorylation via cGKII activation. **A)** Representative immunoblots and a summary graph of normalized phosphorylated tau levels in cultured WT cortical neurons in each condition showing that 1 nM HIV gp120 or 1 nM FIV gp95 treatment significantly increases AT8-positive tau hyperphosphorylation, which is reversed by inhibition of cGKII activity using 1 μM Rp8-Br-PET-cGMPS (RP) (n = number of immunoblots, control (CTRL) = 24, RP = 22, HIV gp120 = 12, HIV gp120 + RP = 10, FIV gp95 = 14, and FIV gp95 + RP = 14, \*\**p* < 0.01 and \*\*\*\**p* < 0.0001, Two-way ANOVA with Tukey test). **B)** Representative immunoblots and a summary graph of normalized phosphorylated tau levels in cultured cGKII KO cortical neurons demonstrating that 1 nM HIV gp120 or 1 nM FIV gp95 treatment has no effect on tau hyperphosphorylation in KO cells (n = number of immunoblots, control (CTRL) = 4, HIV gp120 = 4, and FIV gp95 = 4, One-way ANOVA with Tukey test).

### Viral glycoprotein-induced tau hyperphosphorylation is mediated by cGKII-induced p38 mitogen-activated protein kinase (p38K) activation

Tau can be phosphorylated by various kinases, including p38 mitogen-activated protein kinase (p38K) and glycogen synthase kinase 3 beta (GSK3β) (Billingsley and Kincaid, 1997; Han et al., 2005; Pei et al., 1999; Wang et al., 2013). Among these kinases, p38K is responsible for HIV gp120-induced neuronal death, microglial/macrophage activation, and proinflammatory cytokine production (Kaul and Lipton, 1999; Kaul et al., 2007). Moreover, pharmacological inactivation of p38K in mixed neuronal-glial cultures prevented neuronal death triggered by HIV gp120 (Kaul and Lipton, 1999; Kaul et al., 2007). Importantly, HIV gp120 directly interacts with neurons in the absence of macrophages, and that interaction also results in activation of neuronal p38K (Medders et al., 2010). However, the exact mechanism linking HIV gp120 activation of p38K in both macrophages and neurons currently remains unknown. Previous studies report that activation of p38K requires cGMP-dependent protein kinases in various cell types, including cardiomyocytes, thrombocytes, and fibroblasts (Browning et al., 2000; Kim et al., 2000; Li et al., 2006). We thus hypothesized that HIV gp120 and FIV gp95-induced cGKII stimulation activated p38K and in turn induced tau hyperphosphorylation in neurons. We thus treated DIV 14 mouse cortical neurons with 1 nM HIV gp120 or 1 nM FIV gp95 for 24 hours and determined p38K activation using immunoblots. Notably, HIV gp120 or FIV gp95 treatment significantly increased p38K phosphorylation at threonine 180 and tyrosine 182, an active form of p38K (CTRL, 1.00, HIV gp120, 1.49 ± 0.32, *p* < 0.0001, and FIV gp95, 1.52 ± 0.24, *p* = 0.0012) (**Fig. 4A**). We next examined whether viral glycoprotein-induced p38K activation was dependent on cGKII activation. We added 1 μM RP to DIV 14 cultured neurons treated with 1 nM HIV gp120 or 1 nM FIV gp95 for 24 hours and determined p38K activation using immunoblots as described above. Notably, pharmacological inhibition of cGKII was sufficient to abolish viral glycoprotein-induced p38K activation (HIV gp120 + RP, 0.87 ± 0.29, *p* < 0.0001, and FIV gp95 + RP, 0.92 ± 0.18, *p* = 0.0001) (**Fig. 4A**). However, RP treatment in control neurons had no effect on p38K phosphorylation levels (CTRL + RP, 0.94 ± 0.24, *p* = 0.6229) (**Fig. 4A**). We further used cGKII KO cultured cortical neurons and treated them with 1 nM HIV gp120 or 1 nM FIV gp95 for 24 hours to see if cGKII activation was crucial for viral glycoprotein-induced p38K activation. HIV gp120 or FIV gp95 treatment in KO cells was unable to activate p38K (CTRL, 1.00, HIV gp120, 1.04 ± 0.18, *p* = 0.8773, and FIV gp95, 1.03 ± 0.28, *p* = 0.9462) (**Fig. 4B**), suggesting that neuronal p38K activation by HIV gp120 and FIV gp95 requires cGKII activity.

**Figure 4.**
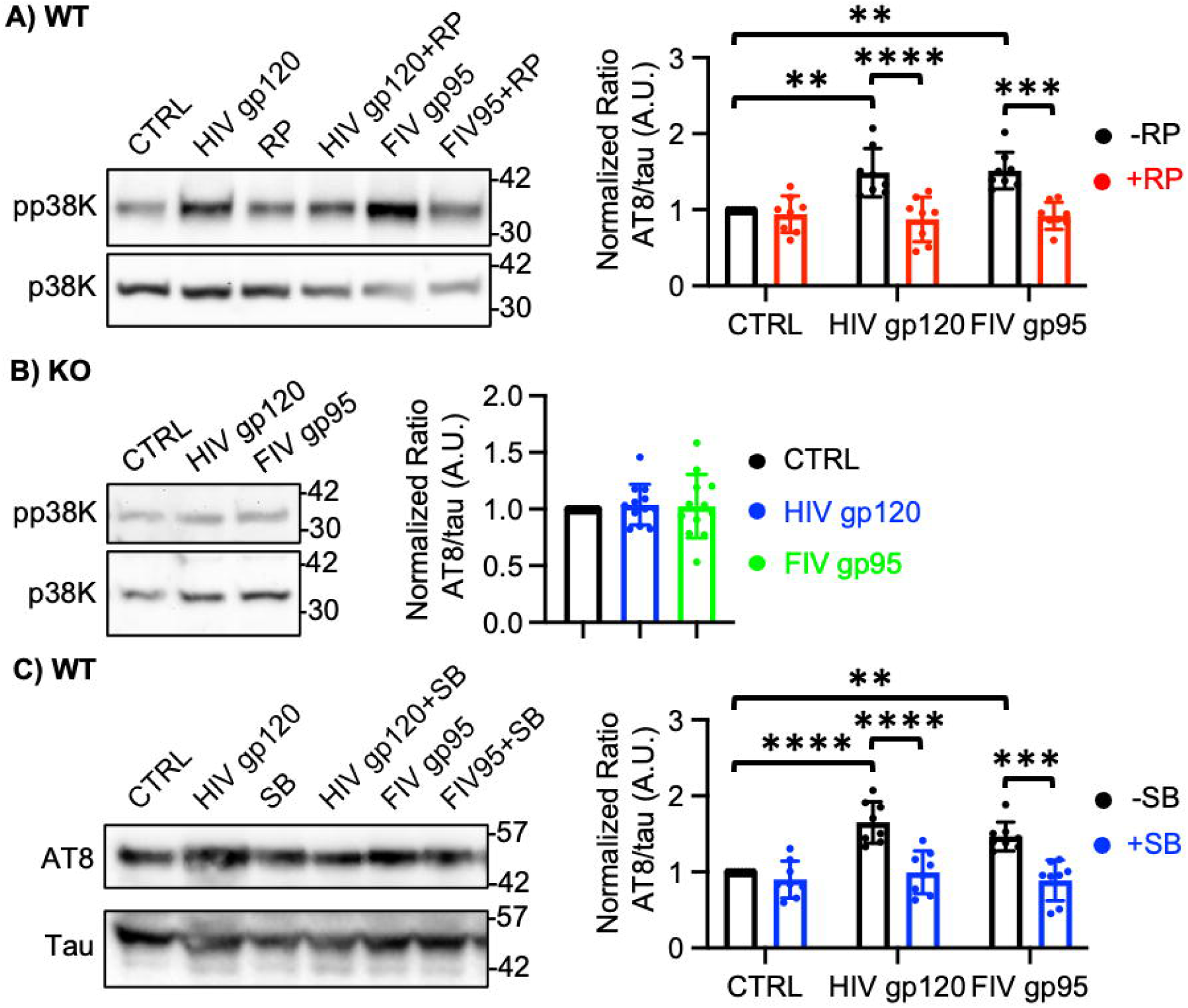
Viral glycoprotein-induced tau hyperphosphorylation is mediated by cGKII-induced p38K activation. **A)** Representative immunoblots and a summary graph of normalized phosphorylated p38K levels in cultured WT cortical neurons in each condition showing that 1 nM HIV gp120 or 1 nM FIV gp95 treatment significantly increases p38K activation, which is reversed by inhibition of cGKII activity using 1 μM Rp8-Br-PET-cGMPS (RP) (n = number of immunoblots, control (CTRL) = 8, RP = 8, HIV gp120 = 8, HIV gp120 + RP = 8, FIV gp95 = 8, and FIV gp95 + RP = 8, \*\**p* < 0.01, \*\*\**p* < 0.001, and \*\*\*\**p* < 0.0001, Two-way ANOVA with Tukey test). **B)** Representative immunoblots and a summary graph of normalized phosphorylated p38K levels in cultured cGKII KO cortical neurons demonstrating that 1 nM HIV gp120 or 1 nM FIV gp95 treatment has no effect on p38K activation in KO cells (n = number of immunoblots, control (CTRL) = 12, HIV gp120 = 12, and FIV gp95 = 12, One-way ANOVA with Tukey test). **C)** Representative immunoblots and a summary graph of normalized phosphorylated tau levels in cultured WT cortical neurons in each condition showing that 1 nM HIV gp120 or 1 nM FIV gp95 treatment significantly increases AT8-positive tau hyperphosphorylation, which is reversed by inhibition of p38K activity using 10 μM SB203580 (SB) (n = number of immunoblots, control (CTRL) = 8, SB = 8, HIV gp120 = 8, HIV gp120 + SB = 8, FIV gp95 = 8, and FIV gp95 + SB = 8, \*\**p* < 0.01, \*\*\**p* < 0.001, and \*\*\*\**p* < 0.0001, Two-way ANOVA with Tukey test).

We next examined whether viral glycoprotein-induced p38K activation was crucial for tau hyperphosphorylation. DIV 14 cultured cortical neurons were exposed to 1 nM HIV gp120 or 1 nM FIV gp95 with 10 μM SB203580 (SB), a p38K inhibitor, for 24 hours. As seen previously (**Fig. 3A**), viral glycoprotein treatment was sufficient to increase phosphorylated PHF tau levels, which was significantly reversed by pharmacological inhibition of p38K activity (CTRL, 1.00, HIV gp120, 1.65 ± 0.27, *p* < 0.0001, FIV gp95, 1.48 ± 0.19, *p* = 0.0028, HIV gp120 + SB, 1.00 ± 0.28, *p* < 0.0001, FIV gp95 + SB, 0.89 ± 0.278, *p* = 0.0001) (**Fig. 4C**). However, SB treatment in control cells had no effect on tau hyperphosphorylation (CTRL + SB, 0.90 ± 0.24, *p* = 0.9526) (**Fig. 4C**). Taken together, we demonstrate that viral glycoproteins, HIV gp120 and FIV gp95, induce tau hyperphosphorylation via cGKII-mediated p38K activation.

### Glutamate treatment is sufficient to induce tau hyperphosphorylation but not mediated by cGKII activation

Using the AT8 antibody, it is shown that glutamate treatment causes a rapid rise in hyperphosphorylated tau protein immunoreactivity in primary neuronal cells (Sindou et al., 1994). It is further suggested that glutamate acts on AMPARs, NMDARs, and metabotropic receptors and can trigger activations of intracellular second messengers leading to tau hyperphosphorylation in cultured cortical neurons (Sindou et al., 1994). However, how glutamate-induced increase in neuronal activity induce tau hyperphosphorylation has not been fully understood. Given that viral glycoprotein-induced neuronal hyperexcitation was sufficient to increase tau hyperphosphorylation via cGKII activation in cultured cortical neurons (**Fig. 1 and 3**), we examined whether cGKII activity was required for glutamate-induced tau hyperphosphorylation. We treated DIV 14 neurons with 1 μM glutamate for 10 min and determined tau hyperphosphorylation using immunoblots with the AT8 antibody as described above and found that glutamate treatment significantly increased hyperphosphorylated tau (CTRL, 1.00 and glutamate, 1.44 ± 0.41, *p* = 0.0300) (**Fig. 5A**). To determine whether such an increase in tau hyperphosphorylation was mediated by cGKII activation, 1 μM RP was added to DIV 14 cultured neurons treated with 1 μM glutamate for 10 min, and we determined tau hyperphosphorylation as described above. Interestingly, RP treatment was unable to reverse glutamate-induced tau hyperphosphorylation (glutamate + RP, 1.83 ± 0.87, *p* = 0.0019) (**Fig. 5A**). To further confirm whether cGKII activation was not required for glutamate-induced tau hyperphosphorylation, we used cultured cGKII KO cortical neurons. DIV 14 cultured cGKII KO neurons were treated with 1 μM glutamate for 10 min, and hyperphosphorylated tau levels were determined by immunoblots. We found that glutamate treatment in KO cells significantly increased tau hyperphosphorylation (CTRL, 1.00 and glutamate, 1.68 ± 0.68, *p* = 0.0345) (**Fig. 5B**). Together, a glutamate-induced increase in neuronal activity significantly elevated hyperphosphorylated tau levels in cultured cortical neurons, which is not mediated by cGKII activation.

**Figure 5.**
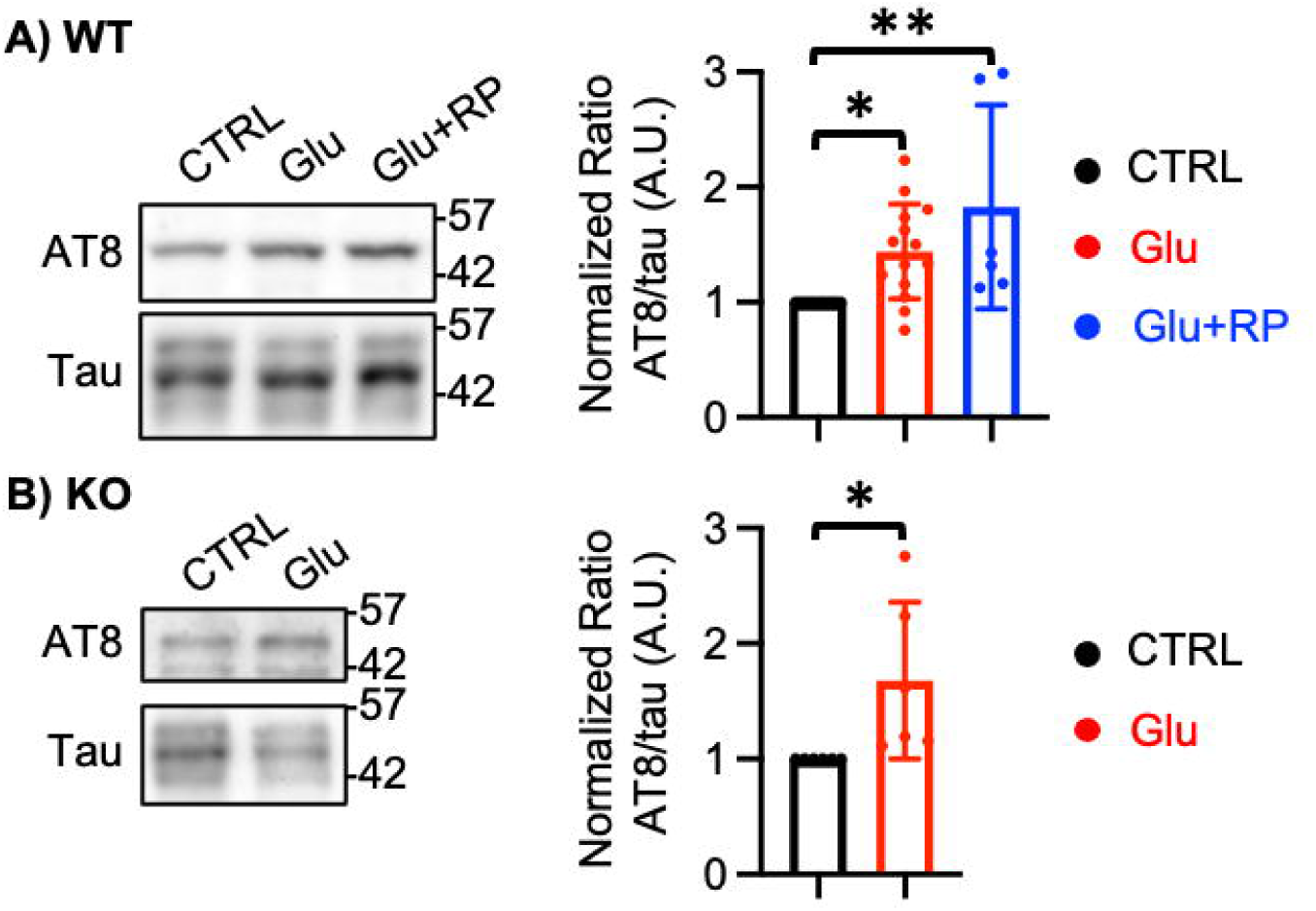
Glutamate treatment is sufficient to induce tau hyperphosphorylation but not mediated by cGKII activation. **A)** Representative immunoblots and a summary graph of normalized phosphorylated tau levels in cultured WT cortical neurons treated with 1 μM glutamate (Glu) or 1 μM glutamate and 1 μM Rp8-Br-PET-cGMPS (Glu + RP) showing that increased neuronal activity significantly increases AT8-positive tau hyperphosphorylation, while cGKII inhibition is unable to prevent such an increase (n = number of immunoblots, control (CTRL) = 14, Glu = 14, and Glu + RP = 6, \**p* < 0.05 and \*\**p* < 0.01, Two-way ANOVA with Tukey test). **B)** Representative immunoblots and a summary graph of normalized phosphorylated tau levels in cultured cGKII KO cortical neurons treated with 1 μM glutamate (Glu) showing that elevated neuronal activity significantly increases tau hyperphosphorylation in cGKII KO cells (n = number of immunoblots, control (CTRL) = 6 and Glu = 6, \**p* < 0.05, unpaired student t-test).

## Discussion

Introduction of cART for HIV has dramatically reduced the number of HIV-related deaths, resulting in more HIV patients are reaching ages when AD-like symptoms appear (Farhadian et al., 2017). Hyperphosphorylated tau, a known AD pathology feature, has been seen in HIV-infected patients’ brains as early as their 30s and further develops as the individuals get older (Anthony et al., 2006). HIV can prematurely age the brains of those with the disease and lead to brain dysfunction, as can AD (Chakradhar, 2018). Given these commonalities, it is possible that HIV could create suitable conditions for the development of AD (Chakradhar, 2018), although the molecular and cellular mechanisms underlying this connection have not been fully investigated (Chakradhar, 2018). Neuronal overexcitation is a putative biological mechanism of HIV-associated AD (Bero et al., 2011; Cirrito et al., 2008; Gibbons et al., 2019; Hinkin et al., 1995; Pooler et al., 2013; Rottenberg et al., 1996; von Giesen et al., 2000; Wu et al., 2016; Yamada et al., 2014). In fact, we have discovered that viral glycoproteins, HIV gp120 and FIV gp95, known neurotoxic viral antigens, stimulate cGKII, which controls AMPA receptor trafficking and hence induces neuronal hyperexcitation (Sztukowski et al., 2018) (**Fig. 6**). Here, our new data using cultured mouse cortical neurons further demonstrate that HIV gp120 and FIV gp95 significantly increase cellular tau pathology, including intracellular hyperphosphorylated tau and extracellular tau release (**Fig. 6**). In addition, we find that this cellular tau pathology is dependent on cGKII activation (**Fig. 6**). Thus, our current work identifies a novel mechanism underlying the link between HIV and AD-related tau pathology relevant to an older population of HIV-infected patients.

**Figure 6.**
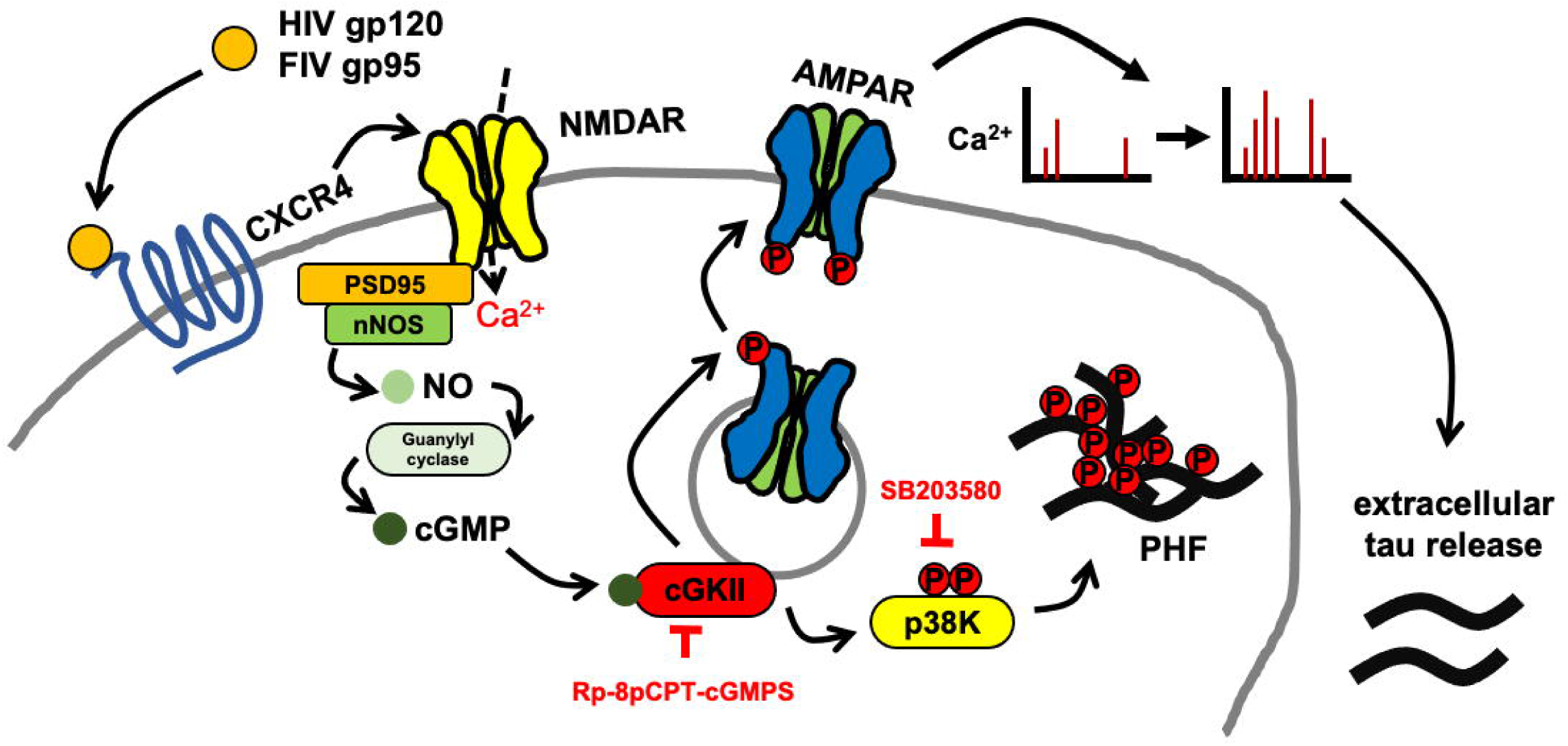
A schematic model of viral glycoprotein-induced cellular tau pathology. FIV gp95 or HIV gp120 interaction with chemokine receptor CXCR4 enhances Ca^2+^-regulating systems through glutamate NMDA receptors (NMDARs). Ca^2+^ fluxes through NMDARs promote the production of nitric oxide (NO) by neuronal nitric oxide synthase (nNOS), which is tethered by the scaffolding protein postsynaptic density 95 (PSD95). NO induces the production of cGMP, in turn activates cGMP-dependent protein kinase II (cGKII) that can phosphorylate a glutamate AMPA receptor (AMPAR) subunit, GluA1, to increase AMPAR synaptic trafficking, which leads to elevation of synaptic excitation. cGKII-induced increase in glutamatergic excitation elicits extracellular tau release, resulting in tau propagation. In addition, cGKII can phosphorylate p38K, leading to tau PHF formation. Therefore, HIV/FIV-induced stimulation of cGKII is critical for AD-related cellular tau pathology. Rp8-Br-PET-cGMPS (RP) and SB203580 (SB) are inhibitors for cGKII and p38K, respectively.

As of now, neither cure nor therapeutic approaches for HIV-associated AD-like pathology are available. Despite the massive investment in AD drugs, there have been more failures than treatment successes (Huang et al., 2019). One of the challenges that the research community has faced is the lack of a viable target for treatment (Saylor et al., 2016). In recent years, a hypothesis implicates the role of neuronal hyperexcitability, hypersynchronous network activity, and aberrant hippocampal network rewiring in memory loss at the early stage of AD (Busche et al., 2012; Busche et al., 2008; Hall et al., 2015; Kazim et al., 2017; Minkeviciene et al., 2009; Siskova et al., 2014). Importantly, brain imaging studies reveal that hypermetabolic signatures of glucose abnormalities, an indication of elevated cerebral neural plasticity, also appear to be an early event in HIV-associated neurocognitive disorder pathogenesis (Hinkin et al., 1995; Rottenberg et al., 1996; von Giesen et al., 2000). However, it remains unclear what are underlying cellular mechanisms of AD and HIV-related neuronal hyperexcitability. We have shown that HIV and FIV glycoprotein-induced neuronal hyperexcitation and cellular tau pathology are mediated by the cGKII-dependent cellular mechanism (**Fig. 6**). This thus suggests that aberrant cGKII stimulation may be a cellular basis for AD and HIV-related neuronal hyperexcitability. Importantly, tau pathology trans-synaptically spreads throughout the brain, which can be stimulated by increased neuronal activity as AD progresses (Braak and Braak, 1991; Kfoury et al., 2012; Sanders et al., 2014; Wu et al., 2016). Indeed, increasing neuronal activity rapidly elevates the steady-state levels of extracellular tau *in vivo* (Sanders et al., 2014; Wu et al., 2016). Moreover, stimulation of excitatory glutamatergic synapses is sufficient to drive tau release (Yamada et al., 2014). Although the mechanism by which tau can be released from neurons is unknown, hyperexcitable neurons in AD and HIV-infected brains may contribute to tau spread. Taken together, abnormal cGKII activation by viral infection can increase neuronal activity via enhancing surface expression of AMPA recetpors, which promotes cellular tau pathology in HIV-positive brains (**Fig. 6**). Therefore, by identifying cGKII as a target of HIV-associated AD pathology, our study completes the pathological pathway and implicates cGKII as a new therapeutic target for limiting HIV-induced AD pathology. Thus, use of cGKII inhibition as a means for neuroprotection may be a novel and innovative approach to this therapeutically challenging disease pathway.

Although multiple lines of evidence suggest that HIV and cART may contribute to AD pathophysiology, little is known about neural mechanisms underpinning HIV-associated neurocognitive disorders and AD in people living with HIV (Brew et al., 2009). In fact, aside from descriptive postmortem studies, different neurodegenerative diseases have been mostly studied in isolation from one another (Brew et al., 2009). Moreover, co-occurrence of neurodegenerative processes complicates treatment development for these diseases (Cohen et al., 2015). One of the major limitations in searching for these disorders is the lack of animal models for studying interactions between neurodegenerative processes in HIV and AD (Chambers et al., 2015).

Thus, new animal models to examine how chronicity and aging affect HIV-induced neuropathology are an important current and future need (Kraft-Terry et al., 2009). The current research provides a novel animal model to address *in vivo* effects of chronic HIV infection on AD-related neural hyperexcitation and tau pathology (tauopathy). There are several *in vitro* and *in vivo* models being used for investigating HIV-induced CNS dysfunction, but these models have limitations in their ability to reproduce current clinical phenotypes (Hatziioannou and Evans, 2012). Furthermore, experimental models of AD-related tauopathy are limited as few species other than humans (Chambers et al., 2015). Although transgenic rodents have been classical animal models of AD, it has been argued that the pathology observed in these animals is different from human AD pathology (Duyckaerts et al., 2008; Frank et al., 2008). Moreover, mice are not naturally susceptible to HIV infection (Fox and Gendelman, 2012). Therefore, additional research is needed to develop better *in vivo* models that consider the effects of chronicity and persistence of viral infection in the CNS (Kraft-Terry et al., 2009). Domestic cats spontaneously develop AD-related tauopathy and are naturally infected with FIV, causing AIDS and neurological complications (Chambers et al., 2015; Head et al., 2005; Janke et al., 1999). Most importantly, we have demonstrated that HIV and FIV glycoproteins employ the same cellular mechanism to induce neuronal hyperexcitation and tau pathology in cultured neurons.

### Based on these features, FIV infection in domestic cats can be an important model for examining the HIV-AD connection

Neuroinflammation plays an important role in AD and possible links to viruses have been proposed (Canet et al., 2018). Microglia are the resident immune cells of the CNS and the major cell types infected by HIV in the brain (Dheen et al., 2007). Neuroinflammatory processes mediated by the activated microglia have been strongly implicated in a number of neurodegenerative diseases including HIV-associated neurocognitive disorders (Gonzalez-Scarano and Baltuch, 1999). Similar to neuronal mechanisms, HIV gp120 interacts with microglial CXCR4 to increase Ca^2+^ activity via enhancing IP3R function, leading to stimulation of inducible NO synthase (iNOS) and subsequent production of NO in microglia (Persichini et al., 2014). Importantly, NO activates cAMP response element-binding protein (CREB) via cGMP-dependent kinase stimulation, leading to upregulation of microglial CD11b expression, an indication of microglial activation during neurodegenerative inflammation (Roy et al., 2006). This suggests that HIV gp120-induced cGKII stimulation in microglia can be critical for their activation. Moreover, HIV gp120 elevates synaptic receptor activity by enhancing the release of proinflammatory cytokines from activated microglia (Kaul et al., 2005; Sillman et al., 2018).

Among those cytokines, tumor necrosis factor alpha (TNFα) induces a rapid increase in GluA1-containing AMPA receptor surface expression and excitatory synaptic strength (Beattie et al., 2002; Stellwagen et al., 2005; Stellwagen and Malenka, 2006). This suggests that microglial cGKII stimulation by viral glycoproteins can be important for activation in microglia, promoting TNFα release, which results in elevation of excitatory synaptic activity, leading to excitotoxicity and tau pathology.

Our cultured neuron system provides cGKII-dependent mechanisms underlying viral glycoprotein-induced neuronal hyperexcitation and cellular tau pathology, including an increase in intracellular hyperphosphorylated tau and extracellular tau release, but our *in vitro* system is unable to produce NFTs. Therefore, further *in vivo* studies are needed to better understand pathophysiology and to develop more effective treatments for AD in HIV infection.

## Methods

### Animals

CD-1 mice (Charles River) were used to produce WT mouse cortical neuron cultures as shown previously (Sathler et al., 2021). cGKII KO animals were maintained as previously described (Kim et al., 2015a; Sztukowski et al., 2018; Tran et al., 2021). We confirmed no cGKII protein expression in cultured KO cortical neurons using immunoblots (**Supplementary Fig. 1**). Colorado State University’s Institutional Animal Care and Use Committee reviewed and approved the animal care and protocol (978).

### Primary cortical neuronal culture

Primary mouse cortical neuron cultures were prepared by a previously described protocol (Kim and Ziff, 2014). Cortices were isolated from postnatal day 0 (P0) CD-1 or cGKII KO mouse brain tissues and digested with 10 U/mL papain (Worthington Biochemical Corp. Lakewood, NJ). Mouse cortical neurons were plated on following poly lysine-coated dishes for each experiment - 24 wells (200,000 cells) for ELISA, glass bottom dishes (500,000 cells) for Ca^2+^ imaging, and 6-cm dishes (2,000,000 cells) for immunoblots. Cells were grown in Neurobasal Medium (Thermo Fisher Scientific) with B27 supplement (Thermo Fisher Scientific), 0.5 mM Glutamax (Thermo Fisher Scientific), and 1% penicillin/streptomycin (Thermo Fisher Scientific).

### GCaMP Ca^2+^ Imaging

DIV 4 neurons were infected with adeno-associated virus (AAV) expressing GCaMP7s (pGP-AAV-syn-jGCaMP7s-WPRE), a gift from Douglas Kim & GENIE Project (Addgene plasmid # 104487; http://n2t.net/addgene:104487; RRID:Addgene_104487) (Dana et al., 2019) for imaging cortical neurons. Neurons were grown in Neurobasal Medium without phenol red (Thermo Fisher Scientific) and with B27 supplement (Thermo Fisher Scientific), 0.5mM Glutamax (Thermo Fisher Scientific), and 1% penicillin/streptomycin (Thermo Fisher Scientific). Glass-bottom dishes were mounted on a temperature-controlled stage on an Olympus IX73 microscope and maintained at 37°C and 5% CO_2_ using a Tokai-Hit heating stage and digital temperature and humidity controller. For GCaMP7s, the images were captured with a 10 ms exposure time and a total of 100 images were obtained with a 1 sec interval. F_min_ was determined as the minimum fluorescence value during the imaging. Total Ca^2+^ activity was obtained by 100 values of ΔF/F_min_ = (F_t_ – F_min_)/F_min_ in each image, and values of ΔF/F_min_ < 0.1 were rejected due to potential photobleaching.

### Reagents

Expression, amplification, and purification of FIV envelope glycoprotein gp95 were performed using previously described methods (de Parseval and Elder, 2001; Sztukowski et al., 2018; Wood et al., 2013). HIV CXCR4-tropic gp120 (IIIB) was obtained through the NIH AIDS Reagent Program, Division of AIDS, NIAID, NIH: HIV-1 IIIB gp120 Recombinant Protein from ImmunoDX, LLC. The following inhibitors were used in this study - 1 μM Rp8-Br-PET-cGMPS (RP) (Thermo Fisher Scientific), 1 μM glutamate (Thermo Fisher Scientific), and 10 μM SB203580 (Thermo Fisher Scientific) for cultured mouse neurons.

### ELISA

Total mouse tau ELISA measurements were carried out using a total mouse tau ELISA kit (Thermo Fisher Scientific) according to the manufacturer’s instruction. DIV 14 cultured mouse cortical neurons were treated with reagents in each experiment, and the conditioned culture medium was collected and centrifuged at 1,000 g for 10 min. Extracellular tau levels were determined in a total tau solid-phase sandwich ELISA with a mouse tau antibody coated plate and mouse tau biotin conjugate. Streptavidin conjugated to horseradish peroxidase was used for assays. All assays were developed using tetramethylbenzidine (TMB) and read on a plate reader at 450 nm. We used recombinant mouse tau as a standard.

### Immunoblots

Immunoblots were performed as described previously (Farooq et al., 2016; Kim et al., 2016; Kim et al., 2018; Shou et al., 2018). The protein concentration in total cell lysates was determined by a BCA protein assay kit (Thermo Fisher Scientific). Equal amounts of protein samples were loaded on 10% glycine-SDS-PAGE gel. The gels were transferred to nitrocellulose membranes. The membranes were blocked (5% powdered milk or 5% BSA) for 1 hour at room temperature, followed by overnight incubation with the primary antibodies at 4°C. The primary antibodies consisted of anti-phosphorylated tau (AT8) (Thermo Fisher Scientific, 1:1000), anti-total tau (Thermo Fisher Scientific, 1:1000), anti-phosphorylated p38K (Cell Signaling Technology, 1:200), anti-p38K (Bioss, 1:1000), anti-cGKII (Kim et al., 2016; Serulle et al., 2007; Tran et al., 2021) (Covance, 1:1000), and anti-actin (Abcam, 1:2000) antibodies. Membranes were subsequently incubated by secondary antibodies for 1 hour at room temperature and developed with Enhanced Chemiluminescence (ECL) (Thermo Fisher Scientific, Waltham, MA). Protein bands were quantified using ImageJ (https://imagej.nih.gov/ij/).

### Statistics

All statistical comparisons were analyzed with the GraphPad Prism 9. Unpaired two-tailed Student’s t-tests were used in single comparisons. For multiple comparisons, we used one-way or two-way ANOVA followed by Tukey test to determine statistical significance. Results were represented as a mean ± SD, and a *p* value < 0.05 was considered statistically significant.

## Acknowledgements

We thank members of the Kim laboratory for their generous support. We also thank Mary Nehring in Dr. Sue VandeWoude’s lab for helping with the ELISA experiments. This work is supported by Student Experiential Learning Grants (SK), COVID-19 Teaching & Research Student Employment Initiative (MJD), and College Research Council Shared Research Program from Colorado State University (SK). This research was also supported by funds from the NIH/NCATS Colorado CTSA Grant (UL1 TR002535), the Boettcher Foundation’s Webb-Waring Biomedical Research Program, and the NIH grant (AG072102-01A1). MJD is a 2019 Boettcher Scholar.

---

**Supplementary Figure 1. Lack of cGKII protein expression in cultured KO cortical neurons.** Representative immunoblots of cGKII protein expression in cultured WT and cGKII KO cortical neurons showing that KO cells have no protein expression. Actin protein expression is used as a loading control.

